# ARHGAP32 as a novel RhoGAP interacting with desmoplakin is required for desmosomal organization and assembly

**DOI:** 10.1101/2023.12.13.571599

**Authors:** Hua Li, Yan Wang, Yinzhen He, Xiayu Liu, Xiufen Duan, Kaiyao Zhou, Gangyun Wu, Wenxiu Ning

## Abstract

Desmosomes are specialized cell-cell junctions that play a critical role in maintaining tissue barrier integrity, particularly in mechanically stressed tissues. The assembly of desmosomes is regulated by the cytoskeleton and its regulators, and desmosomes also function as a central hub for regulating F-actin. However, the specific mechanisms underlying the crosstalk between desmosomes and F-actin, particularly involving RhoGAP or RhoGEF proteins, remain unclear. In our study, we identified that ARHGAP32, a Rho GTPase-activating protein, is located in desmosomes through its interaction with DSP via its GAB2-interacting domain. Using CRISPR-Cas9 gene knockout system, we confirmed that ARHGAP32 is required for proper desmosomal organization, maturation, and length regulation. Notably, the loss of ARHGAP32 resulted in increased formation of F-actin stress fibers and phosphorylation of MYOSIN at T18/S19, indicating enhanced actomyosin contractility. Furthermore, inhibition of ROCK1 kinase activity using Y27632 effectively restored desmosomal organization. Moreover, we demonstrated that the regulation of desmosomes by ARHGAP32 is crucial for maintaining the integrity of epithelial cell sheets. Collectively, our study unveils ARHGAP32 as a RhoGAP present at desmosomes, potentially facilitating the crosstalk between desmosomes and F-actin. Its presence is indispensable for desmosomal assembly and the preservation of epithelial cell sheet integrity by regulating actomyosin contractility.

## Introduction

Cell-cell junctions play a crucial role in the formation and maintenance of tissue barrier integrity (Adil et al., 2021; Garcia et al., 2018; Wu and Sun, 2023). Among the different types of adhesive junctions, desmosomes, acting as “rivets” between keratinocytes, are abundant in tissues that experience considerable mechanical stress, such as the skin (Bharathan et al., 2023; Kowalczyk and Green, 2013). Desmosomal cadherins, including desmogleins (DSG) and desmocollins (DSC), facilitate cell-to-cell connections via extracellular adhesive interactions, while the armadillo protein family members plakoglobin (PKG) and plakophilin (PKP) play a role in forming the intracellular plaque.

The plakin family member desmoplakin (DSP) is proposed to specifically link PKG and PKP to the intermediate filament network (Kowalczyk and Green, 2013). Genetic disorders of desmosomal components cause blistering diseases of the skin, such as pemphigus, as well as other diseases including cardiomyopathies and certain cancers (Broussard et al., 2015; Kottke et al., 2006; Schmidt et al., 2019; Spindler et al., 2023).

The actin cytoskeleton is reported to be involved in the assembly and turnover of desmosomal components (Fuchs et al., 2023; Godsel et al., 2010; Hiermaier et al., 2021; Moch et al., 2022). The relocation of DSP to the desmosome during desmosome assembly can be accelerated by active RhoA, but the maturation of desmosome plaques is hindered by persistent activation of RhoA (Godsel et al., 2010). Overexpressing a catalytic DHPH domain of the RhoA activator ARHGEF11, which enhanced actomyosin contractility, also leads to down-regulation of DSP and DSG1 expression (Ning et al., 2021). Additionally, induction of a microtubule-severing protein, spastin, in differentiated epidermal cells, which also leads to increased actomyosin contractility, causes a significant decrease in DSG1, DSP, and DSC2/3 levels, resulting in reduced desmosome size (Muroyama and Lechler, 2017). Therefore, overload actomyosin contractility caused by increased RhoA should be controlled and limited to facilitate desmosomes assembly. However, little is known how F-actin was controlled during desmosomal turnover and whether there are F-actin inhibitors associated with desmosomes.

On the other hand, desmosomes also function as a signaling hub to influence actin organization (Muller et al., 2021). PKP1 co-localized with F-actin and is essential for cortical actin organization. Lack of desmosomal binding domain of PKP1 induced filopodia and long protrusion (Godsel et al., 2010; Hatzfeld et al., 2000; Keil et al., 2016). PKP2 recruits active RhoA to the cell-cell contact interface, driving actomyosin filament reorganization. Similarly, loss of DSP results in increased in cellular migration and filopodia length as well as aberrant Rac1 signaling (Bendrick et al., 2019; Godsel et al., 2010). In pemphigus, where desmosomes are targeted and disrupted by autoantibody against DSG1 and DSG3, RhoA activity has been observed down-regulated and contribute to the pathogenesis of pemphigus (Jin et al., 2021; Spindler and Waschke, 2011; Waschke et al., 2006). However, it is still unclear whether desmosomes regulate stress fiber formation directly or indirectly through Rho guanine exchange factor (GEF) or a RhoGTPase activating protein (GAP).

In this study, we demonstrated that ARHGAP32 located in desmosomes by interacting with DSP via its GAB2 interacting domain. Depletion of ARHGAP32 impairs desmosomal organization, length, as well as its assembly and maturation. ARHGAP32 inhibition also leads to increased stress fiber formation across the nucleus and increased T18/S19 phosphorylated Myosin, while inhibition of ROCK kinase activity rescues the desmosomal localization. Finally, ARHGAP32 is essential for desmosomes formation in HaCaT 3D equivalent as well as the epithelial cell sheet integrity. In summary, our work highlighted a novel RhoGAP that associated with desmosomes that links desmosome dynamics with stress fiber formation.

## Results

Consistent with our previous findings (Ning et al., 2021), we observed that increasing actomyosin contractility resulted in impaired cortical localization of desmosomal components, including DSP and DSG1 in the mouse skin epidermis (Figures 1A-B), as quantified in Figures 1C-D. These results suggest that a stronger actomyosin contractility is not preferred for desmosomal assembly *in vivo*. Therefore, it is reasonable to speculate there are F-actin inhibitors associated with desmosomes to control actomyosin contractility during the rapid turnover of desmosome (Figure 1E).

**Figure 1.**
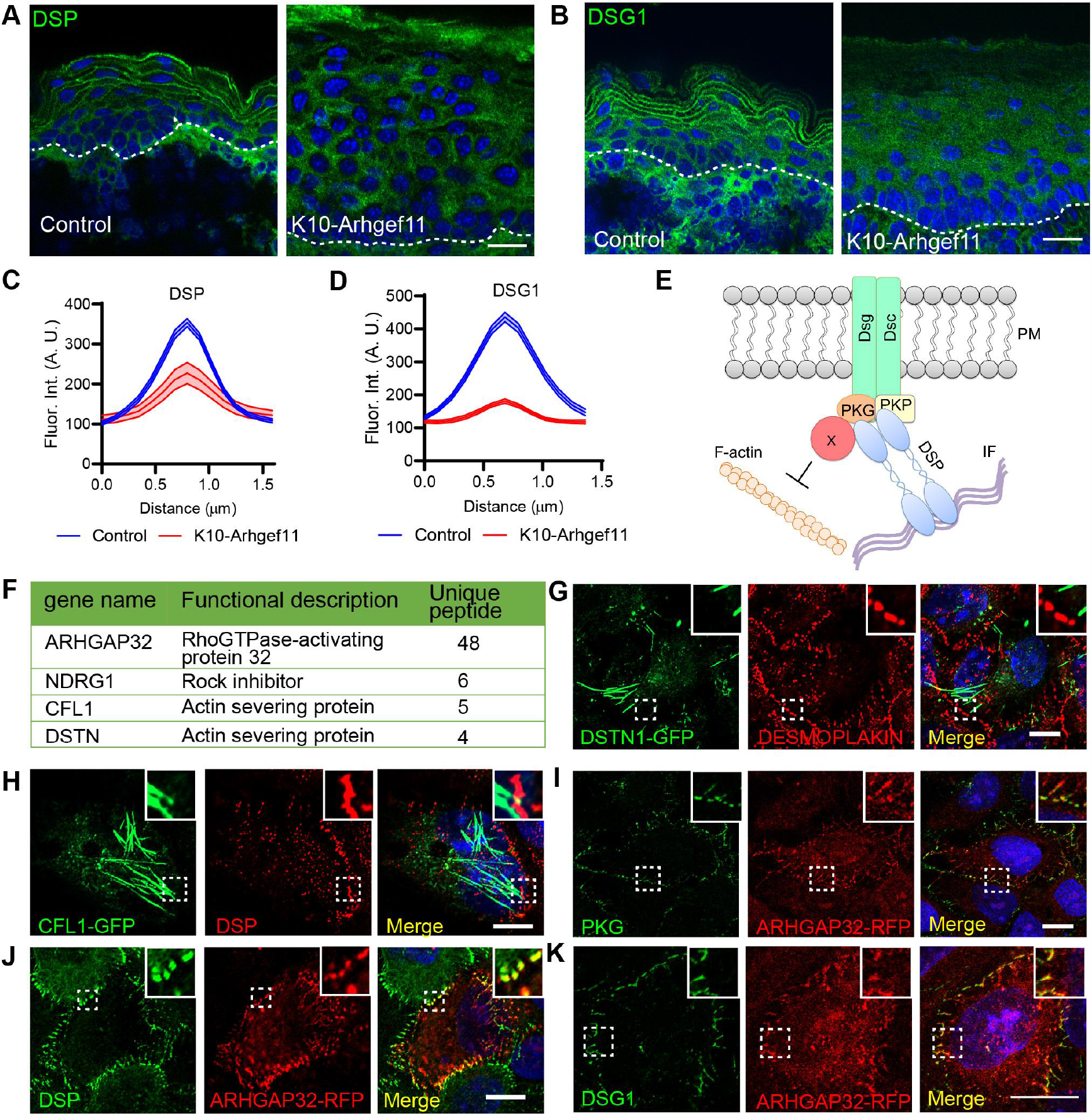
Screening of F-actin inhibitors in Dsp interactomes identified ARHGAP32 co-localized with desmosomes. **A-B**. Immunofluorescence staining of Dsp (A) and Dsg1 (B) in the epidermis of control and dox induced K10rtTA; TREArhgef11 mouse, respectively. **C-D**. Measurement of the fluorescent intensity of Dsp (C) or Dsg1 (D) in control and K10-Arhgef11 mouse epidermis as shown in A-B, respectively. For C, data is mean ± SEM, n=34 for control and n=30 for k10-Arhgef11, p-value <0.0001, two-tailed unpaired t-test. For D, data is mean ± SEM, n=31 for control and n=27 for k10-Arhgef11, p-value <0.0001, two-tailed unpaired t-test. **E**. Diagram of the assumption that an F-actin inhibitor might exist around desmosomes to control F-actin dynamics during desmosomes assembly. **F**. List of F-actin inhibitors and detected relative unique peptide number in DSP interactomes. **G-H**. Co-immunostaining of DSTN1-GFP (G) or CFL1-GFP (H) with DSP in HaCaT cells, respectively. **I-K**. Co-immunostaining of ARHGAP32-RFP with endogenous PKG (**I**), DSP (**J**), and DSG1 (**K**) in HaCaT cells. Scale bars in A and B are 20 μm, while in G-K are 10 μm.

To figure out the possible desmosomes-associated F-actin inhibitors, we screened the proteomic datasets of DSP interactome as published before (Badu-Nkansah and Lechler, 2020), and four candidates that are connected to actin including ARHGAP32, NDRG1, CFL1 and DSTN were selected for further validation (Figure 1F). ARHGAP32 is a GTPase-activating protein (GAP) promoting GTP hydrolysis on RhoA, CDC42, and RAC1 small GTPases (Diring et al., 2019; Okabe et al., 2003). NDRG1 inhibits ROCK1/pMLC2 pathway that modulates stress fiber assembly (Sun et al., 2013). CFL1 and DSTN are actin severing proteins that sever actin filaments (Kanellos and Frame, 2016; Yeoh et al., 2002). The numbers of the unique peptide for these four proteins were listed and indicated ARHGAP32 has a great potential to be associated with desmosomes. We failed to clone the NDRG1 cDNA for the plasmid construction. Moreover, neither DSTN1-GFP nor CFL1-GFP showed co-localization with desmosomal components (Figures 1G-H). Notably, .RHGAP32 was observed co-localized with desmosomal components including PKG, DSP, and DSG1 in HaCaT keratinocytes, respectively (Figures 1I-K). Interestingly, ARHGAP32 co-localized with DSP at cell cortex and as puncta in the cytoplasm (Figure S1). The co-localization of ARHGAP32 with desmosomes was further confirmed in DLD1 epithelial cells (Figures S2A-C).

ARHGAP32 is a large protein consisting of a PX domain, SH3 domain, RHO-GAP domain, GAB2-interacting domain (GAB2-ID), and FYN domain (Figure 2A). To determine how ARHGAP32 localizes at desmosomes, we created truncations that depleted each of the indicated domains of ARHGAP32 and examined their desmosomal localization. Interestingly, the truncations lacking GAB2-ID showed no desmosomal localization, while those containing GAB2-ID still localized at desmosomes (Figures 2B-G). To further confirm the GAB2-ID mediates the desmosomal localization of ARHGAP32, we generated constructions of ΔGAB2-ID-RFP of which the GAB2-ID is deleted of ARHGAP32 and GAB2-ID-RFP with only the GAB2-ID truncations. ΔGAB2-ID-RFP also did not exhibit desmosomal localization (Figure 2H), indicating that the GAB2-ID alone is necessary for the desmosomal localization of ARHGAP32 (Figure 2I). The GAB2-ID mediated desmosomal localization was further confirmed in DLD1 cells (Figures S2D-J). These findings suggest that GAB2-ID is required and essential for the desmosomal localization of ARHGAP32 (Figure 2J).

**Figure 2.**
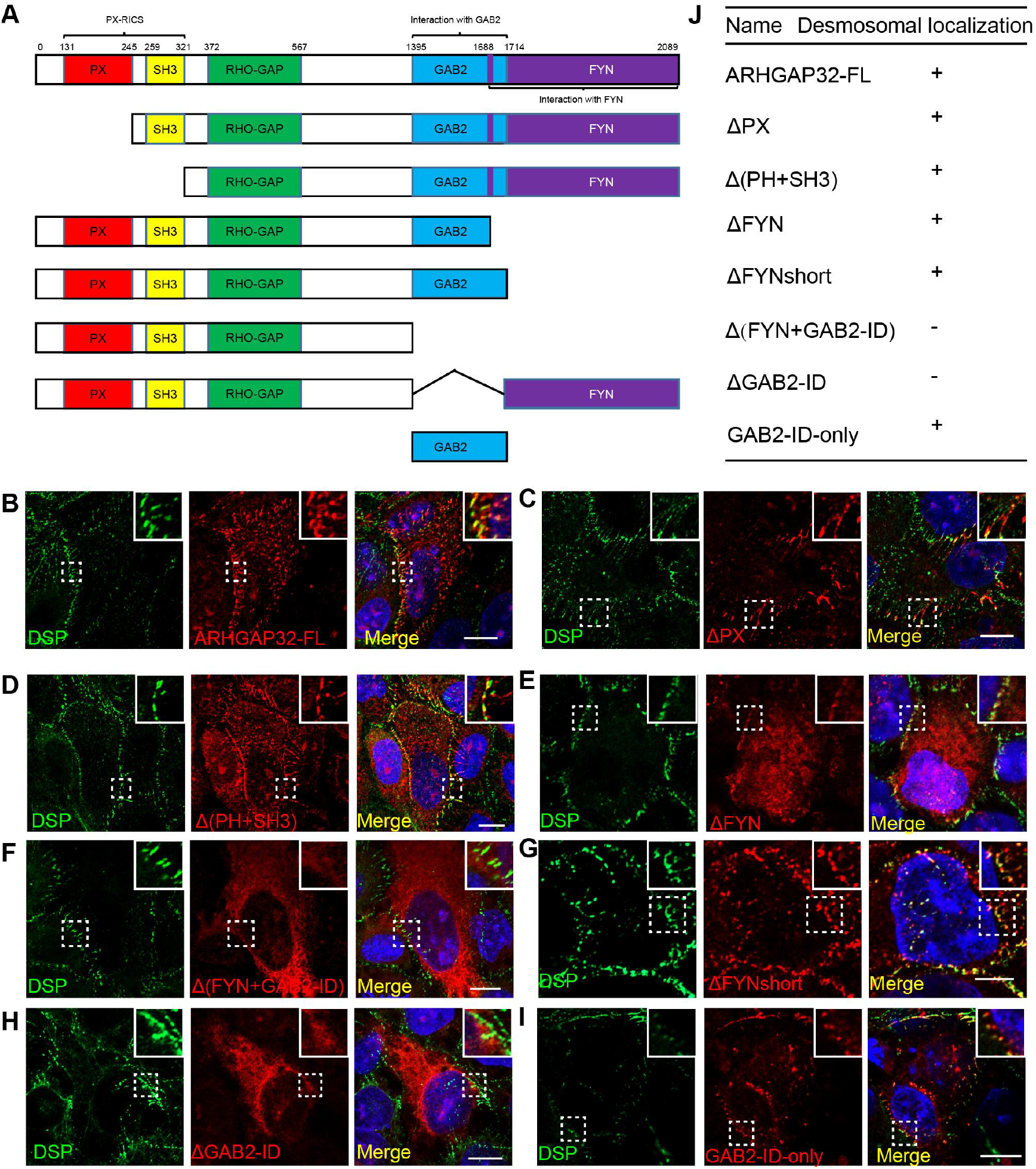
GAB2 interacting domain mediates the desmosomal localization of ARHGAP32. **A**. Diagram of Arhgap32 domain and related truncation forms. The domains include PX domain, SH3 domain, RHO-GAP domain, GAB2 interacting domain (GAB2-ID) and FYN domain. **B-I**. Immunofluorescence staining of endogenous DSP (green) with full-length ARHGAP32-RFP (**B**), ΔPX-ARHGAP32-RFP (**C**), Δ(PH+SH3)-ARHGAP32-RFP (**D**), ΔFYN-ARHGAP32-RFP (**E**), Δ(FYN+GAB-ID)-ARHGAP32-RFP (**F**), ΔFYNshort-ARHGAP32-RFP (do not contain the overlap region in GAB2-ID) (**G**), ΔGAB2-ID-ARHGAP32-RFP (**H**), and GAB2-ID-RFP (**I**), respectively. **J**. Summary of the desmosomal localization of all the truncated mutants of ARHGAP32-RFP. Scale bars are all 10 μm.

To investigate the localization of ARHGAP32 at desmosomes, we initially examined its potential interaction with DSP since the proteomic analysis revealed their proximity. We employed the BiFC technique as a means to validate the protein-protein interactions.

Compared with control transfection of ARHGAP32-VC155 with VN173, the BiFC signal of ARHGAP32-VC155 with DSP-VN173 showed robust fluorescence at the desmosomes (Figure 3A), indicating ARHGAP32 may indeed interact with DSP. Further immunoprecipitation experiment confirmed the interaction between ARHGAP32 and DSP, specifically through the GAB2-ID of ARHGAP32 (Figure 3B-C).

**Figure 3.**
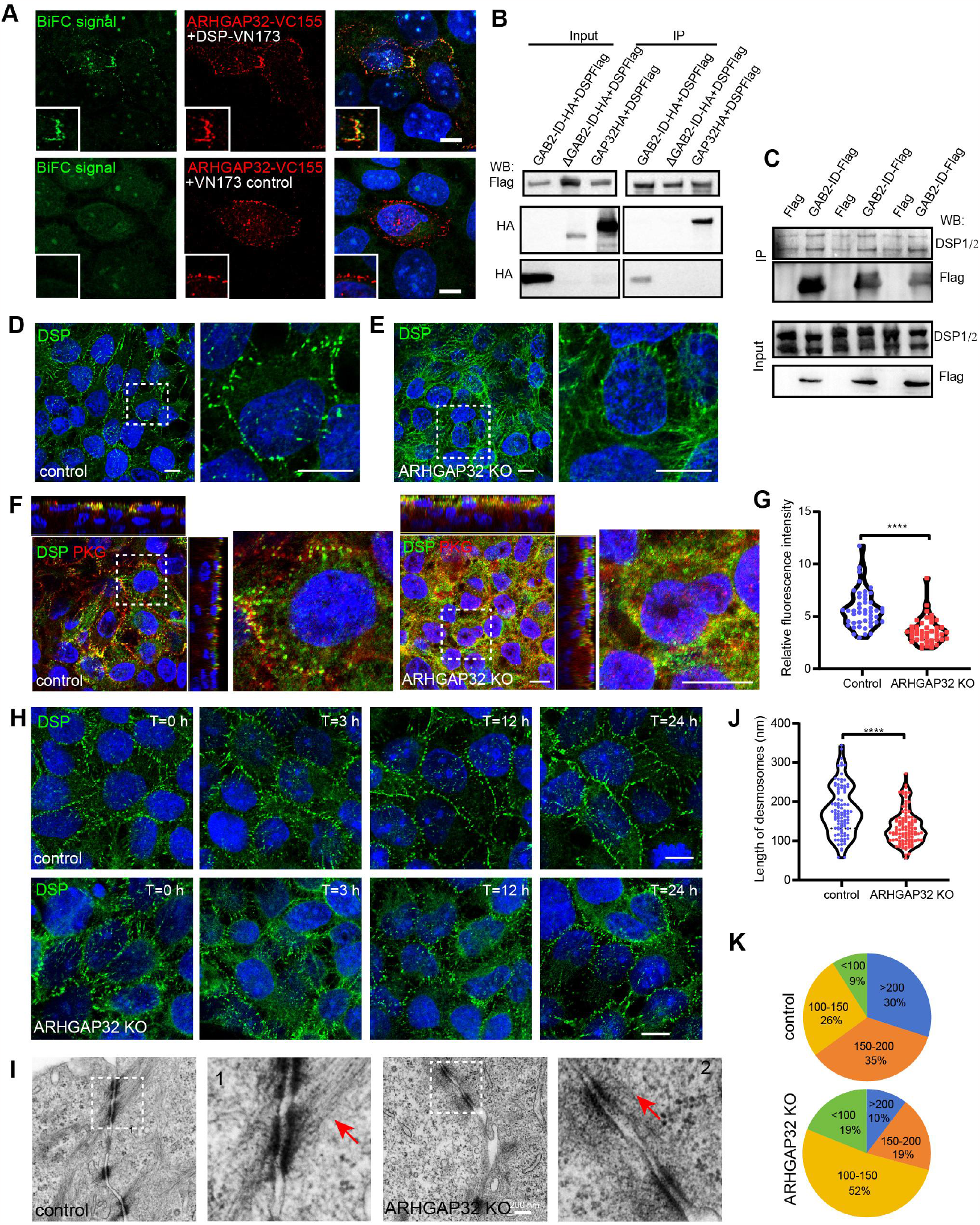
ARHGAP32 interacts with DSP via its GAB2 interacting domain and is required for desmosomal organization and assembly. A. The BiFC signal of ARHGAP32-VC155 with DSP-VN173 or VN173 control in HaCaT cells, respectively. **B**. Immunoprecipitation of DSP-Flag with GAB2-ID-HA, ΔGAB2-ID-HA, or ARHGAP32-HA, respectively in 293T cells. **C**. Immunoprecipitation of GAB2-ID-Flag and Flag control with endogenous DSP in 293T cells. **D-E**. Immunofluorescence staining of DSP in control (**D**) or *ARHGAP32* KO HaCaT cells (**E**), respectively. **F**. Immunofluorescence staining of DSP and PKG in control and *ARHGAP32* KO HaCaT 3D equivalent. **G**. Quantification of the relative fluorescence intensity of desmosomal DSP in control and *ARHGAP32* KO cells as shown in D-E. Data is mean ± SEM, n=47 for control and n=44 for *ARHGAP32* KO cells, p-value <0.0001, two-tailed unpaired t-test. **H**. Immunofluorescence staining of DSP in control and *ARHGAP32* KO HaCaT cells at different time stage after calcium switch. **I**. Transmission Electron Microscopy images of desmosomes in control and *ARHGAP32* KO HaCaT cells. **J**. Quantification of desmosomal length in control and *ARHGAP32* KO cells as shown in I. Data is mean ± SEM, n=100 for control and n=106 for *ARHGAP32* KO cells, p-value <0.0001, two-tailed unpaired t-test. **K**. Percentage of the relative length of desmosomes in control and *ARHGAP32* KO cells as shown in I. Scale bar in I is 200 nm, and the rest are 10 μm.

To investigate whether ARHGAP32 plays a role in regulating desmosomes organization and assembly, we generated stable *ARHGAP32* knockout (KO) HaCaT cells by CRISPR-Cas9 gene editing method, and the exact mutant sites were confirmed by DNA sequencing (Figure S3A). Interestingly, compared to the control, the DSP distribution in *ARHGAP32* KO HaCaT cells exhibited elongated fibers and small puncta within the cells (Figures 3D-E), which was consistently observed in two independent *ARHGAP32* KO cell lines (Figure S3). Additionally, in air-liquid HaCaT 3D equivalent assay, we demonstrated impaired desmosomal localization of DSP and PKG, indicating ARHGAP32 is required for desmosomal assembly in *ex vivo* skin equivalent models (Figures 3F-G). Next, we performed calcium switch assays using control and *ARHGAP32* KO cells (Figure 3H). ARHGAP32 loss delayed desmosomes assembly compared to control cells, as evidenced by the observations at three hours post calcium switch. This indicates that ARHGAP32 is required for efficient desmosomal assembly. Transmission electron microscopy (TEM) analysis of desmosomes in control and *ARHGAP32* KO cells was also conducted (Figure 3I). The TEM images revealed that desmosomes in *ARHGAP32* KO cells were significantly smaller in size (Figure 3J), with the majority ranging from 100-150 nm, while those in control cells ranged from 150-200 nm (Figure 3K). These findings collectively suggest that ARHGAP32 not only plays a role in desmosome organization but is also necessary for desmosomal assembly.

To study the mechanism by which ARHGAP32 regulates desmosomes, we first tested whether the known interacting proteins co-localized with ARHGAP32 at desmosomes. The result showed that GAB1, GAB2 as well as RASA1 did not co-localize with ARHGAP32 at desmosomes (Figures 4A-C). These data further supported that the desmosomal localization of ARHGAP32 is GAB2 protein independent despite GAB2-interacting domain mediate this process. Although CRK has been reported to localize at desmosomes (Badu-Nkansah and Lechler, 2020), we observed weak desmosomal co-localization between CRK and ARHGAP32 (Figure 4D). Interestingly, while RhoA itself did not localize at desmosomes (Figure 4E), ARHGAP32 truncation ΔGAB2-ID-RFP showed strong co-localization with RhoA in the cytoplasm (Figure 4F). This implies that desmosomes may serve as a structure to sequester RhoA activation in the cytoplasm. We further found the F-actin stress fibers were increased after *ARHGAP32* depletion compared with the control cells (Figure 4G), with increased F-actin stress fibers extending across the nucleus (Figure 4H), indicating the increased actomyosin contractility. We next performed the immunofluorescence staining and immunoblotting assay in control and ARHGAP32 depleted cells for the T18/19 phosphorylated MYOSIN which is a hall mark of the increased actomyosin contractility. It further confirmed the increased T18/S19 phosphorylated MYOSIN after ARHGAP32 depletion, consistent with the increased stress fibers (Figures 4I-J). Treatment with ROCK inhibitor Y27632 rescued the desmosomal organization in *ARHGAP32* KO cells (Figures 4K-L). These data indicate that ARHGAP32 is required for desmosomal localization by regulating actomyosin contractility. Furthermore, dispase treatment revealed the increased cell debris after *ARHGAP32* depletion compared to control (Figures 4M-N), meaning ARHGAP32 is required for cell sheet integrity.

**Figure 4.**
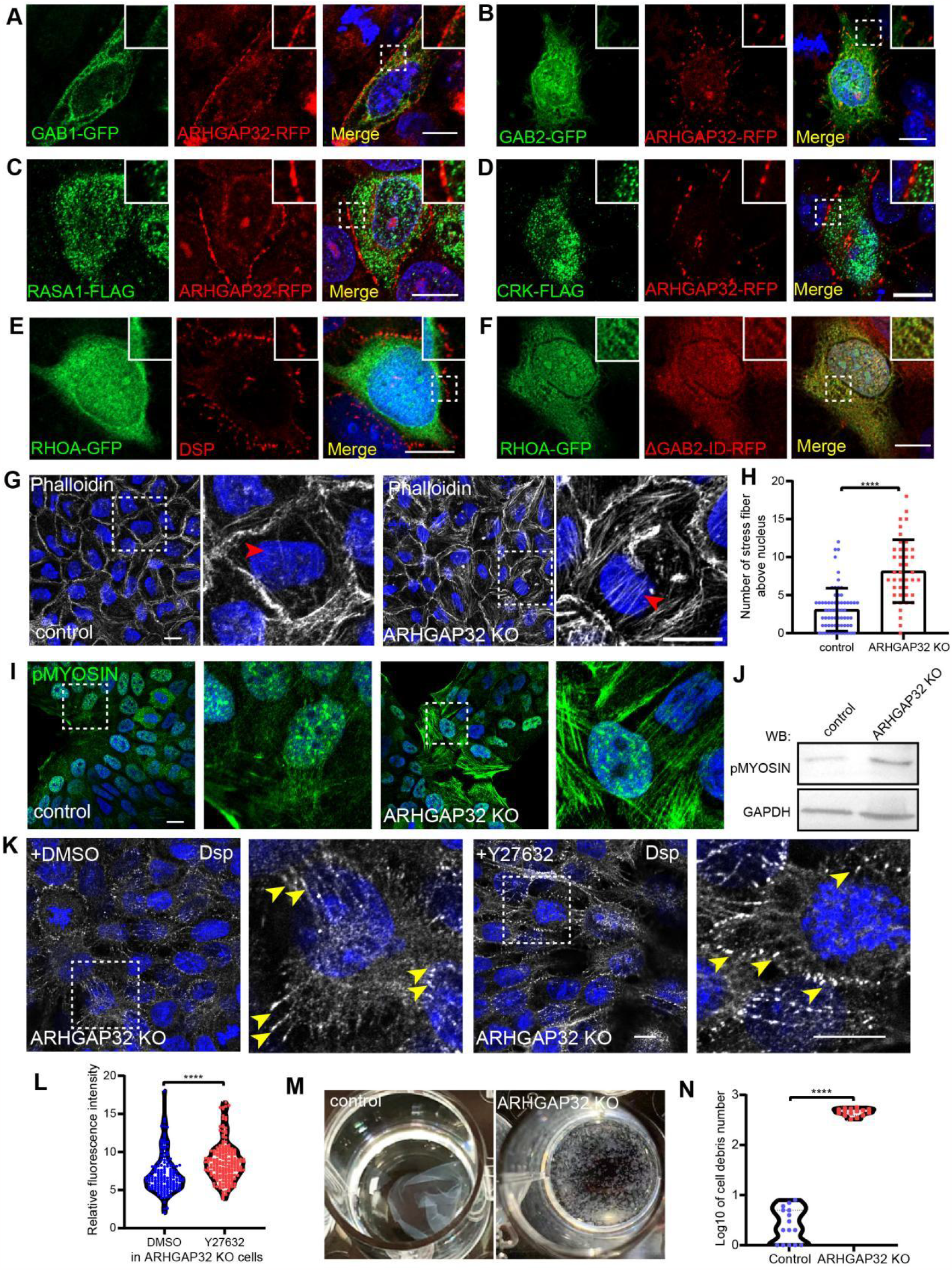
ARHGAP32 depletion affects stress fiber formation and epithelial cell sheet integrity. **A**-**D**. Co-immunostaining of ARHGAP32-RFP with GAB1-GFP (**A**), GAB2-GFP (**B**), RASA1-FLAG (**C**) and CRK-Flag (**D**) in HaCaT cells. **E**. Co-immunostaining of RhoA-GFP with endogenous DSP in HaCaT cells. **F**. Co-immunostaining of RhoA-GFP with ΔGAB2RFP in HaCaT cells. **G**. Immunostaining of F-actin (Phalloidin) in control and *ARHGAP32* KO HaCaT cells. The dashed frame was enlarged at right. Red arrowheads indicate F-actin stress fiber above the cell nucleus. **H**. Quantification of the numbers of stress fiber above cell nucleus related to G. Data is mean ± SEM, n=66 for control and n=42 for *ARHGAP32* KO cells. **I-K**. Immunofluorescence staining (I) and Immunoblotting, (J) of phosphorylated MYOSIN (T18/S19), (K) DSP with DMSO and Y27632 treated control and *ARHGAP32* KO HaCaT cells. Yellow arrowheads in K indicate the desmosomes dots at cell periphery. **L**. Quantification of the relative fluorescence intensity of DSP in DMSO and Y27632 treated *ARHGAP32* KO cells as shown in K. Data is mean ± SEM, n=165 for control and n=196 for *ARHGAP32* KO cells. **M**. Dispase-based-dissociation assay (DDA) in control and *ARHGAP32* KO cells. **N**. The log_10_ of the cell debris number is calculated as shown in M. Data is mean ± SEM, n=15 for control and *ARHGAP32* KO cells. p-value <0.0001, two-tailed unpaired t-test. Scale bars are all 10 μm.

In conclusion, we found ARHGAP32, a RhoGAP, interacts with DSP by its GAB2-ID and it is essential for desmosomal organization as well as assembly likely by regulating actomyosin contractility, which is necessary for preserving cell sheet integrity (Figure S4). In our condition, deletion of *ARHGAP32* increases F-actin stress fiber formation, implying the increased RhoA activity. Consequently, active RhoA accelerates DSP relocation but the persistent activation hinders the desmosome plaque maturation (Godsel et al., 2010). Thus, desmosome locating ARHGAP32 might participate in regulating RhoA activity and F-actin contractility during desmosome assembly and maturation. However, the precise mechanism by which the cytoskeleton regulates desmosomes remains unclear and requires further investigation. Notably, the deletion of GAB2-ID of ARHGAP32 leads to its release into cytoplasm, where it co-localizes with RhoA. This suggests that desmosomes may act as a hub for fine-tuning RhoA activity by sequestering ARHGAP32 through DSP. In pemphigus, auto-antibodies against DSG disrupt desmosomes and affect downstream signaling, including altered RhoA activity, p38 MAPK, and protein kinase C (Bektas et al., 2013; Schmidt et al., 2019). It is possible that in pemphigus, when desmosomal assembly is blocked by the auto-antibody results in the release of ARHGAP32 into the cytoplasm which causes the decreased RhoA activity. Furthermore, we observed co-localization of ARHGAP32 and its GAB2-ID with DSP in the cytoplasm (Figures S1B and S2j). In fibroblasts, ARHGAP32 has been reported to mediate transport of the N-cadherin/β-catenin complex from the endoplasmic reticulum to Golgi (Nakamura et al., 2008). This implies that cytoplasm co-localization of ARHGAP32 with DSP may facilitate the transport of desmosomal component to cell adhesions. However, further investigation is required to fully understand this phenomenon.

## Methods

### Cell culture and transfection

HaCaT cells, DLD1, and HEK293T cells were cultured in DMEM (Gibco, USA) supplemented with 10% fetal bovine serum (Vivacell, China), 100 units/ml penicillin, and 100mg/ml streptomycin (Beyotime, China). The cells were grown in a humidified incubator at 37°C and 5% CO_2_ and passaged at approximately 80-90% confluence. HEK293T cells were transfected with PEI (Polysicences, USA) reagent for virus generation. Stable *ARHGAP32* KO HaCaT cell lines were generated following the protocol by Shalem *et al*. (Shalem et al., 2014) followed by puromycin selection and confirmation *ARHGAP32* mutation through the single colony PCR and sequencing. HEK293T cells for immunoprecipitation assay were transfected using Highgene plus transfection reagent (ABclonal, China). Transient transfections of HaCaT or DLD1 cells were performed using Lipo3000 reagent (Thermo Fisher, USA). In the desmosomes calcium switch experiment, HaCaT keratinocytes were cultured in low calcium medium then switched to high Ca^2+^ (1.5 mM) media. For the actin drug treatment experiment, cells were treated with 10 μM Y-27632 (Glpbio, USA) or DMSO for 2 hours at room temperature before fixation for immunofluorescence staining and imaging.

### Mice

All animal work was approved by Yunnan University. Mice were maintained in a barrier facility with 12-h light/dark cycles. Mouse strains used in this study were obtained and generated as described before.

### Plasmids and primers

CFL1, DSTN full length CDS were PCR amplified from human cDNA and inserted into pEGFP-N1 plasmid by infusion cloning (Vazyme, C112). ARHGAP32 full length CDS and its truncations were also amplified and inserted into pRFP-N1. The pRFP-N1 plasmid was acquired by removing GFP fragment and replaced with a RFP fragment of pEGFP-N1 plasmid. For the bimolecular fluorescence complementation assay, ARHGAP32 and DSP CDS were inserted into pBiFC-VC155 (Addgene #22015) and pBiFCVN173 (Addgene #22010). For immunoprecipitation assay, ARHGAP32 full length CDS and its truncations were PCR amplified and cloned into pCMV-HA plasmid with an HA tag, PCR amplified DSP full length CDS was cloned into pCMV-Tag 2B plasmid with a flag tag. For CRISPR-Cas9 mediated gene knockout mutant construct, the following sgRNA for ARHGAP32 sgRNA1: CGAACTCAACATTCTCATAA, sgRNA2: CCTGACAAGCAATCTGCACG were inserted into lentiCRISPR-V2 (Plasmid #52961) by T4 ligation (NEB, M0202S).

### Immunofluorescence staining

Tissues or cells were immobilized with 4% paraformaldehyde at 37°C for 10 min or with methanol at -20°C for 3 min. After fixation, cells were washed with 1x PBS containing 0.1% triton and then blocked with 1% BSA for 30 min. Then, the samples were incubated with the specified primary antibody for 1 hour at RT, washed 3 times by PBST, and then incubated with secondary antibodies for 30min. After washing with PBST, coverslips or slides were sealed with anti-fluorescence quenching agent (UElandy, China). Frozen blocks were sectioned into 10 µm sections on a cryostat, stained as described above, and mounted on glass slides. The following primary antibodies were used: rabbit anti-DSP (Proteintech, 25318-1-AP), anti-DSG 1 (Proteintech, 24587-1-AP), anti-DSG1 (Santa Cruz, SC-137164), PKG (Santa Cruz, SC-8415), anti-Flag (Sigma, F1804), anti-HA (Invitrogen, 26183), anti-phospho-MYL9-T18/S19 Rabbit (Abclonal, AP0955). The secondary antibodies were used: Phalloidin-Coralite 488 (Proteintech, PF00001), Phalloidin-Coralite 594 (Proteintech, PF00003), Coralite594-conjugate goat anti-rabbit IgG(H+L) (Proteintech, SA00013-4), Coralite594-conjugate goat anti-mouse IgG(H+L) (Proteintech, SA00013-3), Coralite488-conjugate goat anti-rabbit IgG(H+L) (Proteintech, SA00013-2), Coralite488-conjugate goat anti-mouse IgG(H+L), (Proteintech, SA00013-1), DAPI (Meilunbio, MA0127).

### Dispase-based dissociation assay (DDA)

The DDA experiment was modified as previously described [40]. Briefly, the human keratinocytes in 24 well plates were grown to confluent in 10% FBS medium. Then the cells were washed twice with phosphate buffered saline and incubated with 2.4 U/mL dispase (Sigma, USA) at 37°C for 1 hour. The floating monolayer was rotated in a shaker (300 rpm) for 5-10 min. A photograph of the fragment was taken with the camera and the cell counter function was used to determine the amount of fragment using the ImageJ software.

### Immunoprecipitation assay

HEK293T cells were transfected with indicated flag-tagged plasmids for 36 hours with 6 cm plates and the cell lysate were extracted using RIPA buffer. Then the immunoprecipitation procedure was carried out using an anti-flag affinity gel kit (Beyotime, China). The eluted proteins were then analyzed by western blotting. The primary and secondary antibodies used were as below: anti-Flag (Sigma, F1804, USA), anti-HA (Invitrogen, 26183, USA), anti-phospho-MYL9-T18/S19 Rabbit (Abclonal, AP0955, China), and anti-DSP (Proteintech, 25318-1-AP, China), Horse Anti-mouse IgG, HRP-linked Antibody (CST, 7076S, USA), Goat Anti-rabbit IgG, HRP-linked Antibody (CST, 7074S, USA)

### Transmission electron microscopy

Cell samples were washed with PBS and fixed with 2.5% glutaraldehyde on ice for 2 h, then washed in PBS 3 times. Cells were then fixed with 1% osmium tetroxide for 1 h and washed with distilled water 3 times, then stained with 2% uranyl acetate for 30 min, and wash 3 times with distilled water. Samples were then dehydrated in 50%, 70%, 80%, 90% and 100% ethanol for 2 min respectively, then were infiltrated in 50%, 2/3, 3/4 epoxy resins for 30 min, finally were infiltrated in 100% resin overnight at room temperature. On the second day, samples were embedded in fresh resin for 1 hour and labeled, then polymerized at 60°C for 48 h. After polymerization, the samples were section using an EM UC7 ultramicrotome, then stained with 2% uranyl acetate for 10 min, and washed with distilled water for 10 min, then stained with 0.2% lead citrate for 5 min, and finally washed with distilled water for 10 min. After the samples were naturally dried, imaging was performed by transmission electron microscope (HITACHI, HT7800).

### Quantifications

Images were analyzed and quantified using FIJI software. The fluorescence intensities of cortical DSP and DSG1 were measured by drawing lines across cell junctions, and their profiles were analyzed and plotted for significance and relative p-value based on the highest fluorescence intensity at cell junctions. The relative fluorescence intensity of DSP and PKG at desmosomes was calculated by inviding the highest fluorescence intensity at desmosomes by their average fluorescence intensity in the cytoplasm. Desmosomes length was analyzed by measuring the length of dense desmosomes in SEM images. The number of stress fiber above the nucleus was quantified. DDA assay quantified the numbers of debris until control cell sheet were shaked off from plates after treated with dispase. All experiments were performed at least three times independently. Statistical significance was determined by a two-tailed paired or unpaired Student’s t test, with asterisks denoting significance (ns = not significant, *, p < 0.05; **, p < 0.01, ***, p < 0.001, ****, p < 0.0001). Where no significance is indicated, p values were > 0.05. All statistical analysis was performed using Microsoft Excel.

### Imaging

Images were acquired using Zen software (Zeiss) on a Zeiss AxioImager Z1 microscope with Apotome 2 attachment at either 20X/0.8 objective or 63X/1.4 oil objective, or acquired using Zeiss LSM 800 confocal with Airyscan using 63x oil objective at around 1% laser intensity and master gain at 800v.

## Acknowledgements

We are grateful to Professor Jianwei Sun from Yunnan University for sources of HEK293T and DLD1 cell lines. We thank core facilities and animal center of Yunnan University for imaging and mice care.

## Author contributions

Conceptualization: WX.N., H.L.; Methodology: Y.W., YZ.H., H.L.; Software: WX.N., Y.W., YZ.H., XY.L.; Validation: XY.L., XF.D., KY.Z., GY.W.; Formal analysis: WX.N., H.L., Y.W., YZ.H.; Investigation: Y.W., YZ.H., XY.L., XF.D., KY.Z., GY.W., H.L.; Resources: WX.N.; Writing -original draft: WX.N., H.L.; Writing-review & editing: WX.N., H.L., XY.L., YZ.H.; Visualization: WX.N., H.L., Y.W., YZ.H.; Supervision: WX.N., H.L.; Project administration: WX.N.; Funding acquisition: WX.N., H.L.

---

**Figure S1.**
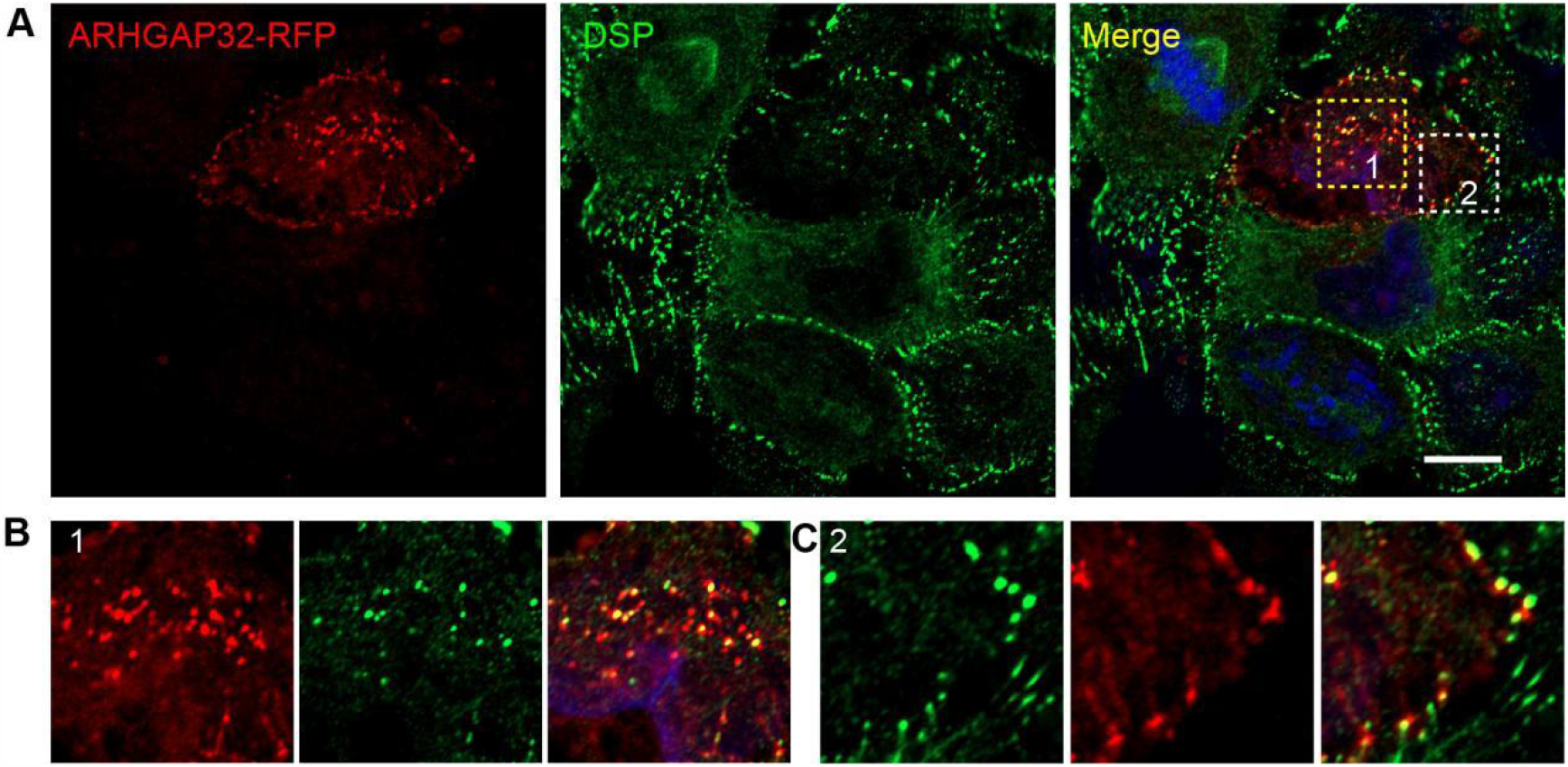
ARHGAP32-RFP co-localizes with DSP both at desmosomes and cytoplasm. **A**. Co-immunostaining of ARHGAP32-RFP with DSP in HaCaT cells. **B**. Co-immunostaining of ARHGAP32-RFP with DSP in the cytoplasm as puncta in HaCaT cells as shown in frame 1 in A. **C**. Co-immunostaining of ARHGAP32-RFP with DSP in the desmosomes as shown in frame 2 in A. Scale bars are all 10 μm.

**Figure S2.**
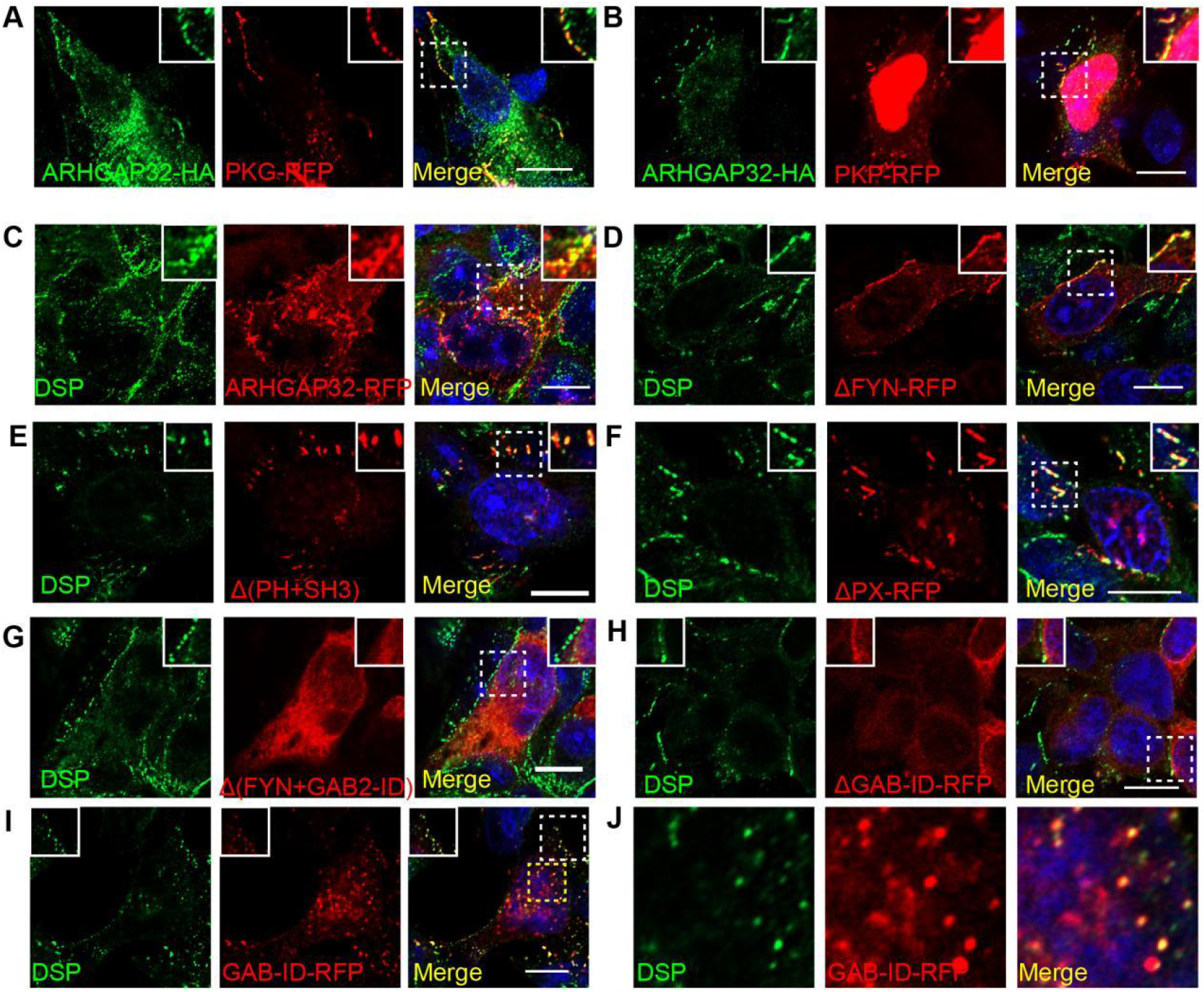
ARHGAP32-RFP co-localizes with DSP at desmosomes in DLD1 cells. **A-B**. Co-immunofluorescence staining of ARHGAP32-HA (green) with PKG-RFP (A), and PKP-RFP (B) in DLD1 cells. **C-J**. Co-immunofluorescence staining of DSP with ARHGAP32-RFP (C), ARHGAP32 truncations of ΔFYN-RFP (D), Δ(PH+SH3)-RFP (E), ΔPX-RFP (F), Δ(FYN+GAB2-ID)-RFP (G), ΔGAB2-ID-RFP (H), GAB2-ID-RFP (I) in DLD1 cells. **J**. Co-localization of ARHGAP32-GAB2-ID-RFP with DSP in the cytoplasm as puncta as shown in the yellow frame in I. Scale bars are all 10 μm.

**Figure S3.**
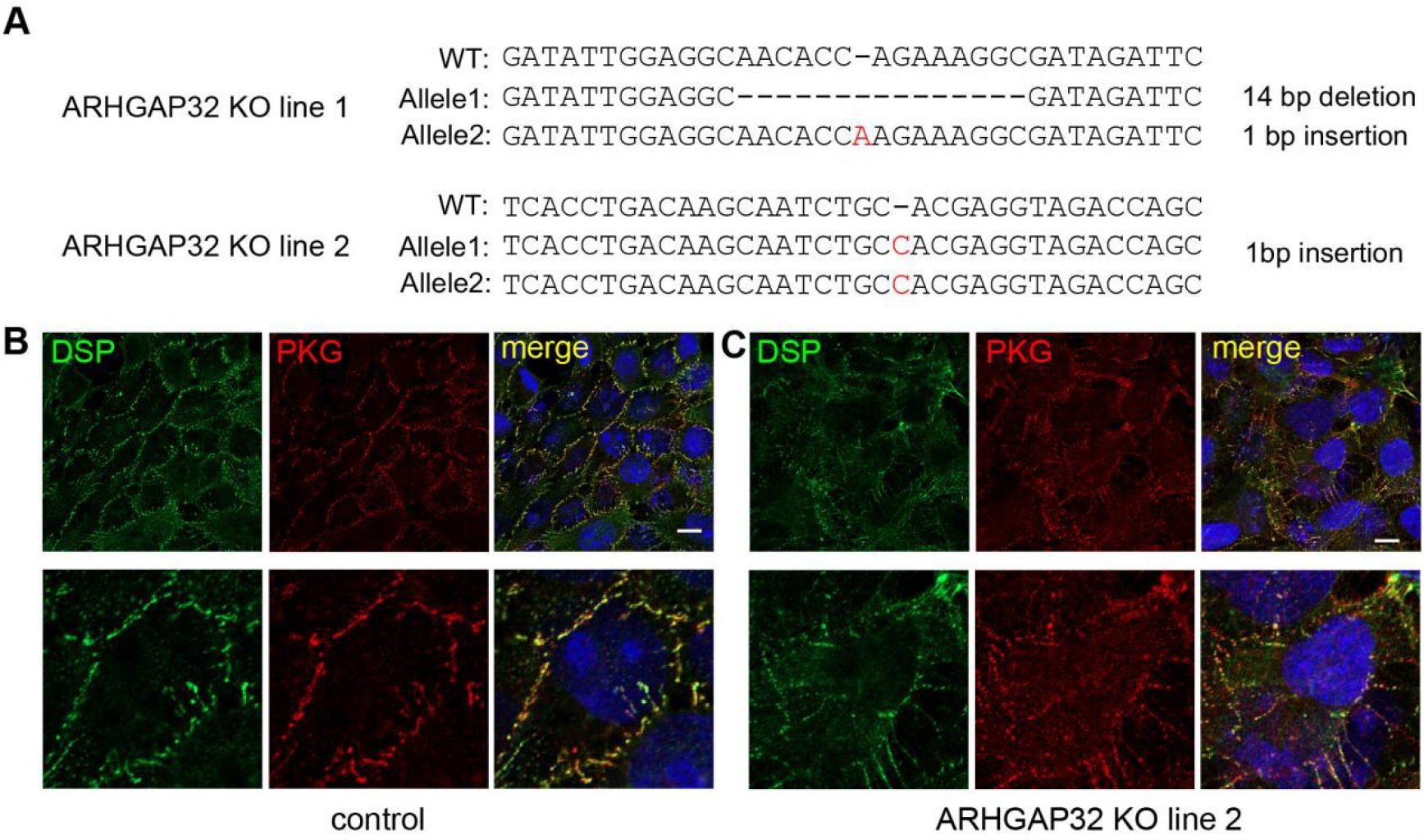
ARHGAP32 is essential for desmosomes organization. **A**. Mutation site of *ARHGAP32* sequence after CRISPR-Cas9 knock out. For *ARHGAP32* KO line 1 as shown in Figure 3D, there is a 14 bp deletion in allele 1, and a 1 bp insertion in allele 2. For *ARHGAP32* KO line2 as shown in Figure S3B, there is a 1 bp insertion in 2 alleles. **B**. Co-immunofluorescence staining of DSP and PKG in *ARHGAP32* KO HaCaT cell line 2. Scales bars are 10 μm.

**Figure S4.**
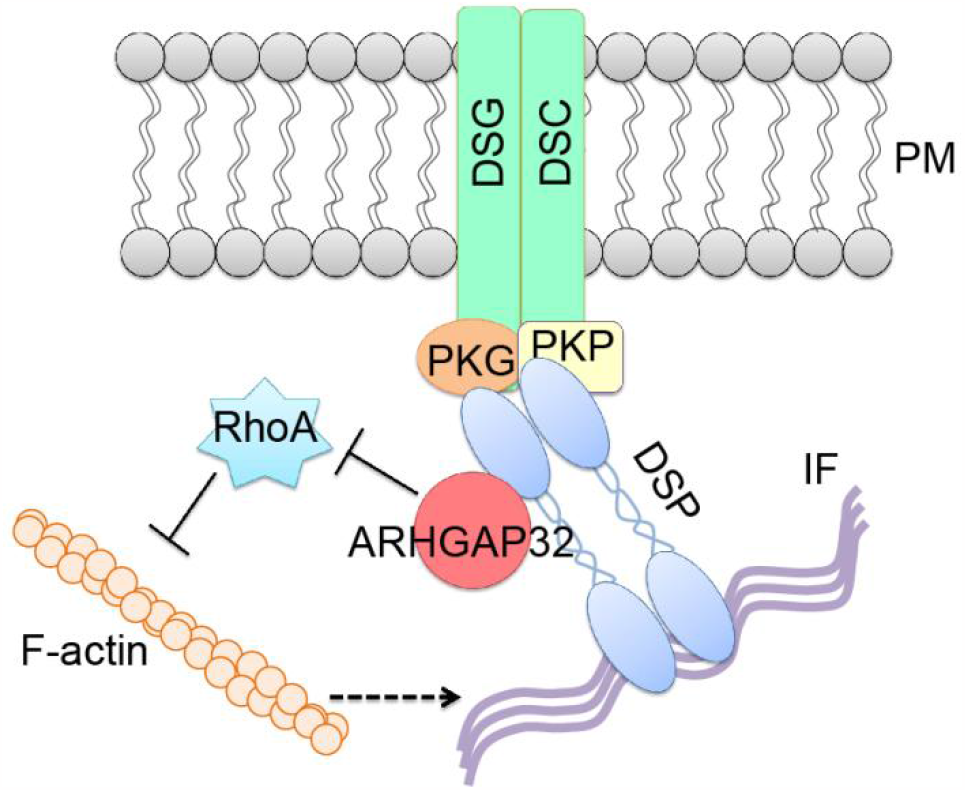
Model of ARHGAP32 in the regulation of desmosomes. ARHGAP32 interacts with DSP in the desmosomes, while it helps to limit RhoA activity to inhibit F-actin stress fiber formation whose enhancement has been shown to affect desmosomes assembly.

